# Differential activation of G-protein-mediated signalling by synthetic cannabinoid receptor agonists

**DOI:** 10.1101/850651

**Authors:** Shivani Sachdev, Samuel D. Banister, Marina Santiago, Chris Bladen, Michael Kassiou, Mark Connor

**Author notes:** **Corresponding author**: Mark Connor. Department of Biomedical Sciences, Macquarie University, NSW, Australia.

## Abstract

Synthetic cannabinoid receptor agonists (SCRAs) are new psychoactive substances associated with acute intoxication and even death. However, the molecular mechanisms through which SCRAs may exert their toxic effects remain unclear - including the potential differential activation of G protein subtypes by CB1, a major target of SCRA. We measured CB1-mediated activation of Gα_s_ and Gα_i/o_ proteins by SCRAs by examining stimulation (PTX-treated) as well as inhibition (non-PTX treated) of forskolin-induced cAMP accumulation in HEK cells stably expressing CB1. Real-time measurements of stimulation and inhibition of cAMP levels were made using a BRET biosensor. We found that the maximum concentration of SCRAs tested (10 μM), increased cAMP levels 12 to 45% above that produced by forskolin alone, while the phytocannabinoid THC did not significantly alter cAMP levels in PTX-treated HEK-CB1 cells. All SCRAs had greater potency to inhibit of forskolin-induced cAMP levels than to stimulate cAMP levels. The rank order of potencies for SCRA stimulation of cAMP (Gα_s_) was PB-22 > 5F-MDMB-PICA > JWH-018 > AB-FUBINACA > XLR-11. By contrast, the potency of SCRAs for inhibition of cAMP (Gα_i/o_) was 5F-MDMB-PICA > AB-FUBINACA > PB-22 > JWH-018 > XLR-11. The different rank order of potency of the SCRAs to stimulate Gα_s_-like signalling compared to Gα_i/o_ signalling suggests differences in G protein preference between SCRAs. Understanding the apparent differences among these drugs may contribute to unravelling their complex effects in humans.

## INTRODUCTION

The use of synthetic cannabinoid receptor agonist (SCRA) new psychoactive substances (NPS) is associated with significant morbidity and mortality compared to use of Δ^9^-tetrahydrocannabinol (THC), the main psychoactive ingredient of cannabis [1, 2]. SCRAs are linked to wide range of toxic effects including seizures, agitation, hypertension, cardiotoxicity, kidney damage, and sometimes death [3, 4]. There has been a rapid increase in the number of structurally diverse SCRAs since 2010, with little known about their pharmacology and toxicology at time of identification [5]. The constant evolution of SCRA structures occurs in response to legislative restriction and development of urine drug screens for existing compounds [6, 7, 8]. A time-series of seizures (by tonnage) of NPS reported to United Nations Office on Drug and Crime [9] showed that the SCRAs dominated the synthetic NPS market over the period 2011-2017.

Many SCRAs are agonists at cannabinoid type-1 and type-2 receptors (CB1 and CB2, respectively [10]; with the psychoactive effects attributed to the activation of CB1 [11]. We have previously described the *in vitro* quantitative measurement of SCRA efficacy at CB1, where all SCRAs tested showed between 20-300 fold greater agonist activity at CB1 compared to THC [12]. Cannabinoid receptors mediate downstream signalling predominantly through the Gα_i/o_ protein family [13]; however, under some circumstances, CB1 can also stimulate adenylyl cyclase (AC) through Gα_s_-proteins [14, 15, 16]. For example, blockade of canonical CB1-Gα_i_ pathway with pertussis toxin (PTX) or sequestration of CB1-Gα_i_ protein in the primary striatal rat neurons on co-expression with D2 results in an augmentation of cyclic adenosine monophosphate (cAMP) levels by cannabinoids, suggesting CB1 couples to Gα_s_ [14, 15]. A recent study characterized the relationship between CB1 receptor expression and signalling, and showed that at very high receptor expression levels, the effect of CB1 activation on cAMP signalling was stimulatory, a phenotype that was reversed by systematic pharmacological knockdown at the receptor level [17]. The idea that certain SCRAs may preferentially activate different CB1 Gα subtypes is not unprecedented [18, 19, 20]; in a study by Costain and colleagues [21], AB-CHMINACA elicited an elevation in cAMP levels in both the absence and presence of forskolin in human embryonic kidney (HEK) cells transiently expressing CB1, suggesting an AB-CHMINACA-specific CB1-mediated activation of Gα_s_ signalling.

The mechanism(s) through which SCRAs exert different behavioural and physiological effects remains unclear, and which pathways modulated by CB1 activation mediate the specific pharmacological effects of SCRAs is also unknown. Similarly, the question of whether these pathways are activated in a quantitatively or qualitatively similar way by SCRAs and THC is only beginning to be addressed [22]. Finally, the question of whether SCRA activity at non-cannabinoid receptors is also important for their pharmacological effects is very much open [23, 24, 25]. With more than 250 SCRAs identified in the NPS market [9], elucidation of the differential molecular mechanisms by which these compounds can exert distinct pharmacology, including their signalling via CB1, is essential for understanding their adverse effects. This study examined whether SCRAs that are representative of structural classes confirmed in patients admitted to emergency departments with presumed SCRA toxicity stimulate Gα_s_-like cAMP signalling via CB1. We measured the SCRA-mediated stimulation as well as inhibition of forskolin induced cAMP accumulation in HEK cells stably expressing CB1. We have observed SCRA-specific CB1-dependent activation of the two signalling pathways, but THC only coupled to inhibition, not stimulation of cAMP. While AB-CHIMINACA, previously identified as having a unique profile among SCRAs for elevating cAMP, appeared to signal, in part, through non-CB1 mechanisms.

## METHODS

### CB1 receptor transfection and cell culture

HEK 293 FlpIn cells with homogeneous G protein-gated inwardly rectifying K^+^ (GIRK4) channel expression (the construction of these cells by Grimsey and colleagues will be described elsewhere) were co-transfected with pcDNA5/FRT construct encoding haemagglutinin (HA)-tagged human CB1 receptor cDNA and pOG44 (Flp recombinase plasmid) using transfection reagent Fugene HD (Promega) as previously described for AtT-20 pituitary tumour cells [26]. Cells stably expressing the CB1 receptor were cultured in Dulbecco’s Modified Eagle Media (Thermo Fischer Scientific, Waltham, MA, USA) supplemented with 10% fetal bovine serum (FBS, FBS, Sigma-Aldrich, St. Louis, MO, USA), 100 units ml^−1^ penicillin, 100 μg ml^−1^ streptomycin (Thermo Fischer Scientific, Waltham, MA, USA), 400 μg ml^−1^ G418 (GIRK4 selection antibiotic) and 100 μg ml^−1^ hygromycin (CB1 selection antibiotic) up to passage 5 (selection phase). Hygromycin concentration was reduced to 80 μg ml^−1^ beyond passage 5 (maintenance phase). Cells were grown in 75 cm^2^ flask at 37 °C/5 % CO_2_ and passaged at 80% confluency as required. Assays were carried out on cells up to 25 passages.

### Assay for cAMP measurement

Intracellular cAMP levels were measured using pcDNA3L-His-CAMYEL plasmid, which encodes the cAMP sensor YFP-Epac-RLuc (CAMYEL) as outlined in [27, 28]. Cells were detached from the flask using trypsin/EDTA (Sigma-Aldrich), and resuspended in DMEM supplemented with 10% FBS, 100 units ml^−1^ penicillin, and 100 μg ml^−1^ streptomycin. Cells were seeded in 10 cm dishes at a density of 7,000,000 such that they would be 60-70% confluent the next day. On the following day, the cells were transiently transfected with 5 μg of pcDNA3L-His-CAMYEL plasmid using the linear polyethylenimine (PEI, m.w. 25 kDa) (Polysciences, Warrington, PA, USA). The PEI/DNA complex mixture was sequentially added to the cells at the ratio of 1:6, and cells were incubated in 5% CO2 at 37 °C. Approximately 24 hours after transfection, the cells were then detached from the dish and the pellet was resuspended in Leibovitz’s (L-15 -Thermo Fischer Scientific, Waltham, MA, USA) media supplemented with 1% FBS, 100 units ml^−1^ penicillin, 100 μg ml^−1^ streptomycin and 15 mM glucose. In the experiments with pertussis toxin (PTX) to irreversibly uncouple Gα_i_ proteins, the cells were resuspended in the media containing 200 ng ml^−1^ PTX. The PTX-treated and control (non-PTX treated) cells were plated at a density of 100,000 cells per well in poly D-lysine (Sigma-Aldrich) coated, white wall, clear bottomed 96-well microplates. Cells were incubated overnight at 37 °C in ambient CO_2_.

The day after plating, forskolin (FSK, an activator of AC) was prepared in HBSS composed of (mM) NaCl 145, HEPES 22, Na_2_HPO_4_ 0.338, NaHCO_3_ 4.17, KH_2_PO_4_ 0.441, MgSO_4_ 0.407, MgCl_2_ 0.493, CaCl_2_ 1.26, glucose 5.56 (pH 7.4, osmolarity 315 ± 15), and supplemented with 0.1% BSA. All the drugs used for the series of real-time measurements of stimulation and inhibition of cAMP levels were made in 3 μM of forskolin immediately before the assay. The concentration of DMSO (0.10-0.13%) was kept constant for all experiments, however this limited the maximum drug concentration that could be tested. Coelenterazine H substrate (NanoLight Technologies, Pinetop, AZ, USA) was added to a final concentration of 5 μM to the cells, and incubated for 5 min after which 10 μl of (10X) drug was added to each well to obtain the desired concentration. A vehicle (HBSS plus DMSO alone) was included in each column of 96-well microplate and routinely subtracted from the measurements. The PTX-treated and control cells were compared side by side. Luminescence was measured using PHERAstar plate reader (BMG Labtech, Germany) at 37 °C. The cells signalling was measured at an emission wavelength of 475 nm and 535 nm simultaneously, and the readings were made every 40 s for approximately 20 min. A concentration response curve (CRC) for CP55940 and WIN55212-2 inhibition of cAMP accumulation were performed for each experimental replicate as a reference standard (Figure 1).

**Figure 1.**
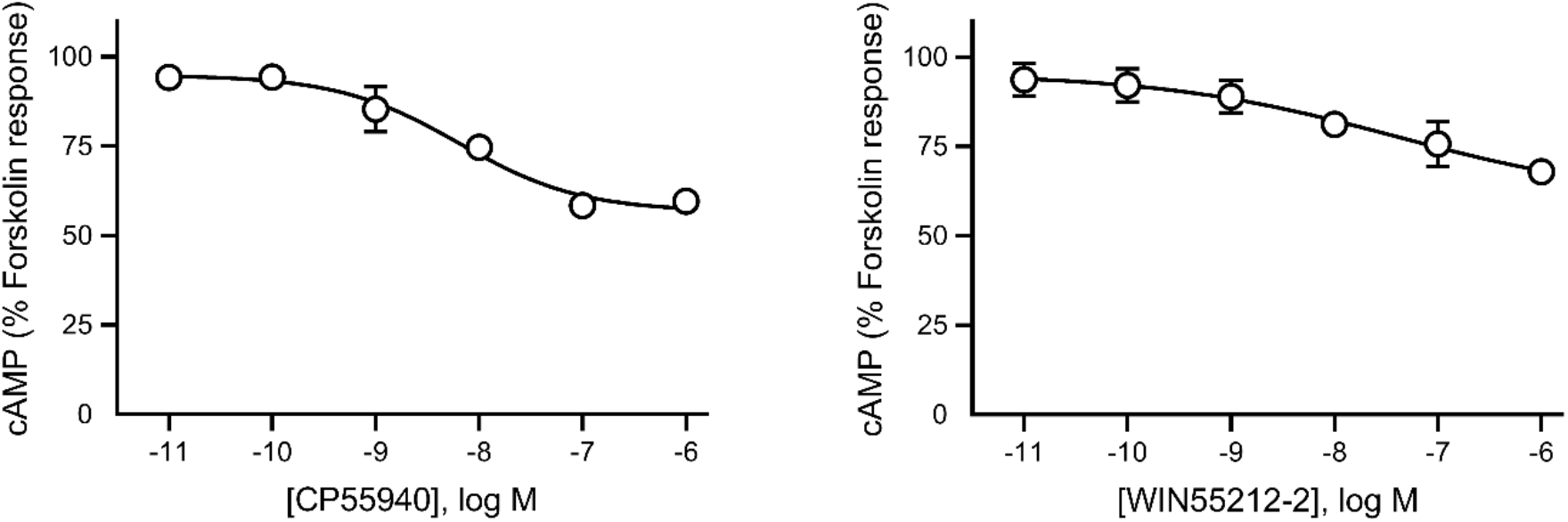
Concentration response curve (CRC) for CP55940 and WIN55212-2. Treatment with CP55940 or WIN55212-2 produced a concentration-dependent inhibition of forskolin-mediated cAMP production in HEK 293-CB1. Area under the curve analysis for CP55,940 or WIN55212-2 in the presence of 3 μM forskolin. Data were normalized to forskolin (100%) and vehicle (0%), and plotted as mean ± SEM for at least 5 independent experiments performed in duplicate.

## Data Analysis

Raw data are presented as inverse bioluminescence resonance energy transfer (BRET) ratio of emission at 475 nm/535 nm, such that an increase in ratio corresponds with increase in cAMP production. Area under curve (AUC) analysis was performed in GraphPad PRISM (Graph Pad Software Inc., San Diego, CA). Data were normalized to forskolin (set as 100%) over a 20 min period for each experiment. Concentration response curves were obtained by fitting four-parameter non-linear regression curves in PRISM to derive EC_50_ and E_MAX_. All final datasets passed the Shapiro-Wilk test for normality. Unless otherwise stated, the data represent mean ± SEM of at least 5 independent experiments, each conducted in duplicate. The differences between groups were tested using unpaired Student’s t-test (PRISM). Statistical significance is defined as P < 0.05.

## Materials

CP55940, WIN55212-2, 2-arachidonoylglycerol (2-AG), CUMYL-4CN-BINACA, and SR141716 were purchased from Cayman Chemical (Ann Arbor, MI, USA), Δ^9^-tetrahydrocannabinol (THC) was from THC Pharm GmbH (Frankfurt, Germany) and was a kind gift from the Lambert Initiative for Cannabis Therapeutics (University of Sydney). PTX was from HelloBio (Bristol, UK), and forskolin was from Ascent Scientific Ltd. All the SCRAs, unless otherwise stated, were synthesized by Dr. Samuel D. Banister in the lab of Professor Michael Kassiou at Sydney University (Sydney, NSW, Australia). Chemical structure of SCRAs can be found elsewhere [12]. All the SCRAs were prepared in DMSO and stored in aliquots of 30 mM in −30 °C until needed.

## RESULTS

### Real-time cAMP BRET measurement of the G_αs_ mediated signalling of SCRAs

Using the CAMYEL assay, we measured the effect of seventeen cannabinoids (10 μM each) on the forskolin-stimulated cellular cAMP levels in HEK-CB1 cells following pre-treatment with PTX. All the SCRAs produced an increase in cAMP levels above that produced by forskolin alone (100%). Examples of raw traces are shown for some SCRAs (Figure 2), note that the stimulation of cAMP by SCRAs in presence of forskolin and PTX plateaued approximately after 12 min, and maintained at that level for the entire course of the assay (20 min). The effects of SCRAs tested ranged from 12 to 45% increase in signal relative to forskolin alone. Most of the SCRAs had approximately 1.5 times higher effect than CP55940 (19%) or WIN55212-2 (18%), except for JWH-018, UR-144, AM-2201, and CUMYL-4CN-BINACA, which showed similar or lower effect (Figure 2). AB-FUBINACA had up to 2.5 times higher effect than CP55940. In PTX treated cells, the endocannabinoid 2-AG (10 μM) produced an increase in forskolin-stimulated cAMP levels approximately twice that of CP55940, while the phytocannabinoid THC did not significantly alter cAMP levels in the presence of forskolin (compared to forskolin alone Figure 2, P > 0.05).

**Figure 2.**
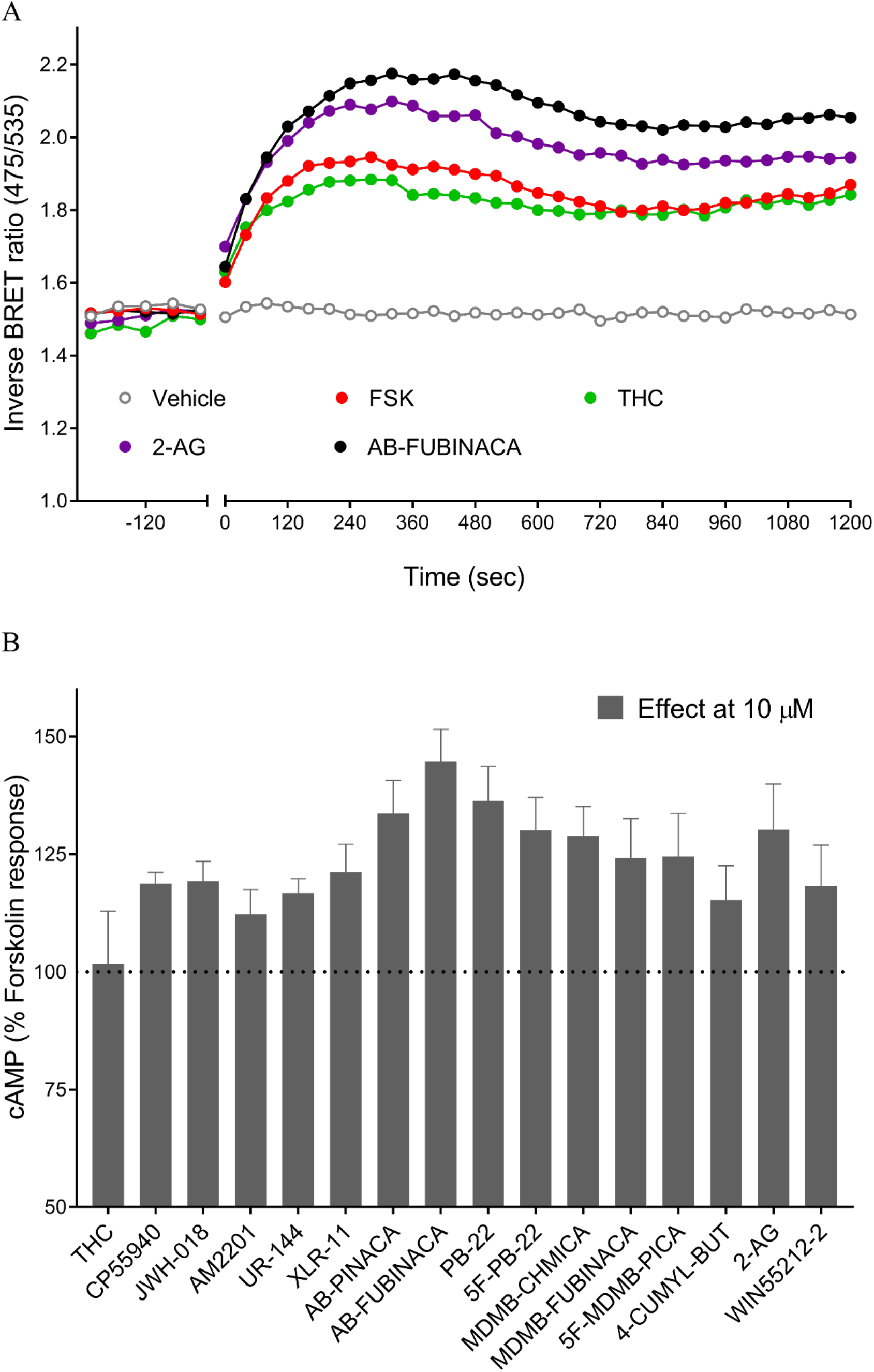
Gα_s_ mediated signalling of SCRAs. **A.** Representative data for real-time measurement of stimulation of cAMP levels by 10 μM of cannabinoids (THC, 2-arachidinoylglycerol, and AB-FUBINACA) in HEK cells expressing CB1 receptors, an increase in inverse BRET ratio (emission at 475/535 nm) corresponds to an increase in cAMP. **B.** A bar chart summarising the cAMP signalling peaks for seventeen cannabinoids showing an increase in cAMP levels above that of forskolin alone (100%). Graphs show mean ± SEM for at least 5 independent experiments performed in duplicate.

### Differential SCRAs-induced stimulation and inhibition of cAMP signalling in HEK-CB1

To assess whether there was any evidence of preferential coupling to Gα_i/o_ over Gα_s_ among SCRAs, we assessed the pharmacological activity (EC_50_ and E_MAX_) of a selection of the most prevalent SCRAs (JWH-018, PB-22, AB-FUBINACA, XLR-11, and 5F-MDMB-PICA), to stimulate and inhibit cAMP in HEK-CB1 cells. All SCRAs tested activated CB1 via Gα_i/o_ (inhibitory, non-PTX treated), and Gα_s_ (stimulatory, PTX-treated) in a concentration dependent manner (Figure 3). As previously reported [29], treatment with CP55940 and WIN55212-2 produced an immediate concentration dependent inhibition of forskolin-mediated cAMP production (*p*EC_50_ CP55940 8.2 ± 0.3, *p*EC_50_ WIN55212-2 7.4 ± 0.2). All SCRAs had greater potency (0.62-16 nM) for inhibition of forskolin-induced cAMP levels in non-PTX treated HEK cells compared to their potency to stimulate cAMP levels. The activation of CB1-Gα_s_ by SCRAs showed a wide variation in E_MAX_ values, with significant differences in efficacies proceeding AB-FUBINACA (most efficacious) > PB-22 ≈ XLR-11 ≈ 5F-MDMB-PICA > JWH-018, whereas all the SCRAs were similarly effective at inhibiting cAMP production (Table 1). The first SCRA to be identified in spice, JWH-018, caused partial (13% increase over forskolin alone) activation of Gα_s_ pathway, but produced greater inhibition of the forskolin-induced cAMP response (64% of forskolin response). Whereas other SCRAs tested in the present study induce moderate activation of Gα_s_ pathway (26-44% relative to forskolin) compared to their activity at Gα_i/o_ inhibitory pathway. The rank order of potencies for SCRAs for inhibition of cAMP (Gα_i/o_) is 5F-MDMB-PICA > AB-FUBINACA > PB-22 > JWH-018 > XLR-11. By contrast, the potency of SCRAs for stimulation of cAMP (Gα_s_) is PB-22 > 5F-MDMB-PICA > JWH-018 > AB-FUBINACA > XLR-11. The most efficacious SCRA at Gα_s_-pathway (AB-FUBINACA) was roughly 350 times less potent at Gα_s_ than the Gα_i/o_-pathway, while JWH-018 was only14 times less potent. XLR-11 had much lower potency compared to all the other SCRAs for both Gα_s_-and Gα_i/o_-pathway.

**Table 1.** Comparison of pharmacological activity (EC_50_ and E_MAX_) of SCRAs-induced stimulation (*G*_*s*_ (+PTX)) and inhibition (*G*_*i*_ (−PTX)) of cAMP signalling in HEK-CB1 cells. The selectivity is expressed as the ratio of *G*_*s*_ (+PTX) EC_50_ to *G*_*i*_ (−PTX) EC_50_.

| Compound | $G_i$ (-PTX) | | $G_s$ (+PTX) | | $G_i$ (-PTX)<br>selectivity |
| --- | --- | --- | --- | --- | --- |
| | p$EC_{50} \pm$<br>SEM<br>($EC_{50}$, nM) | $E_{max}$ (% of<br>FSK) ±<br>SEM | p$EC_{50} \pm$<br>SEM<br>($EC_{50}$, nM) | $E_{max}$ (% of<br>FSK) ±<br>SEM | |
| CP55940 | $8.2 \pm 0.3$<br>(6.4) | $57 \pm 5$ | - | - | - |
| WIN55212-2 | $7.4 \pm 2$<br>(40) | $60 \pm 9$ | - | - | - |
| JWH-018 | $7.8 \pm 0.3$<br>(16) | $64 \pm 4$ | $6.7 \pm 0.7$<br>(221) | $113 \pm 4$ | 14 |
| XLR-11 | $7.2 \pm 0.2$<br>(63) | $62 \pm 3$ | $5.2 \pm 0.8$<br>(6490) | $127 \pm 16$ | 103 |
| PB-22 | $8.6 \pm 0.2$<br>(2.5) | $64 \pm 3$ | $7.2 \pm 0.5$<br>(69) | $131 \pm 5$ | 28 |
| AB-FUBINACA | $9.0 \pm 0.2$<br>(1.1) | $61 \pm 2$ | $6.4 \pm 0.5$<br>(383) | $144 \pm 12$ | 348 |
| 5F-MDMB-PICA | $9.2 \pm 0.2$<br>(0.62) | $59 \pm 4$ | $7.1 \pm 0.4$<br>(85) | $126 \pm 5$ | 137 |

**Figure 3.**
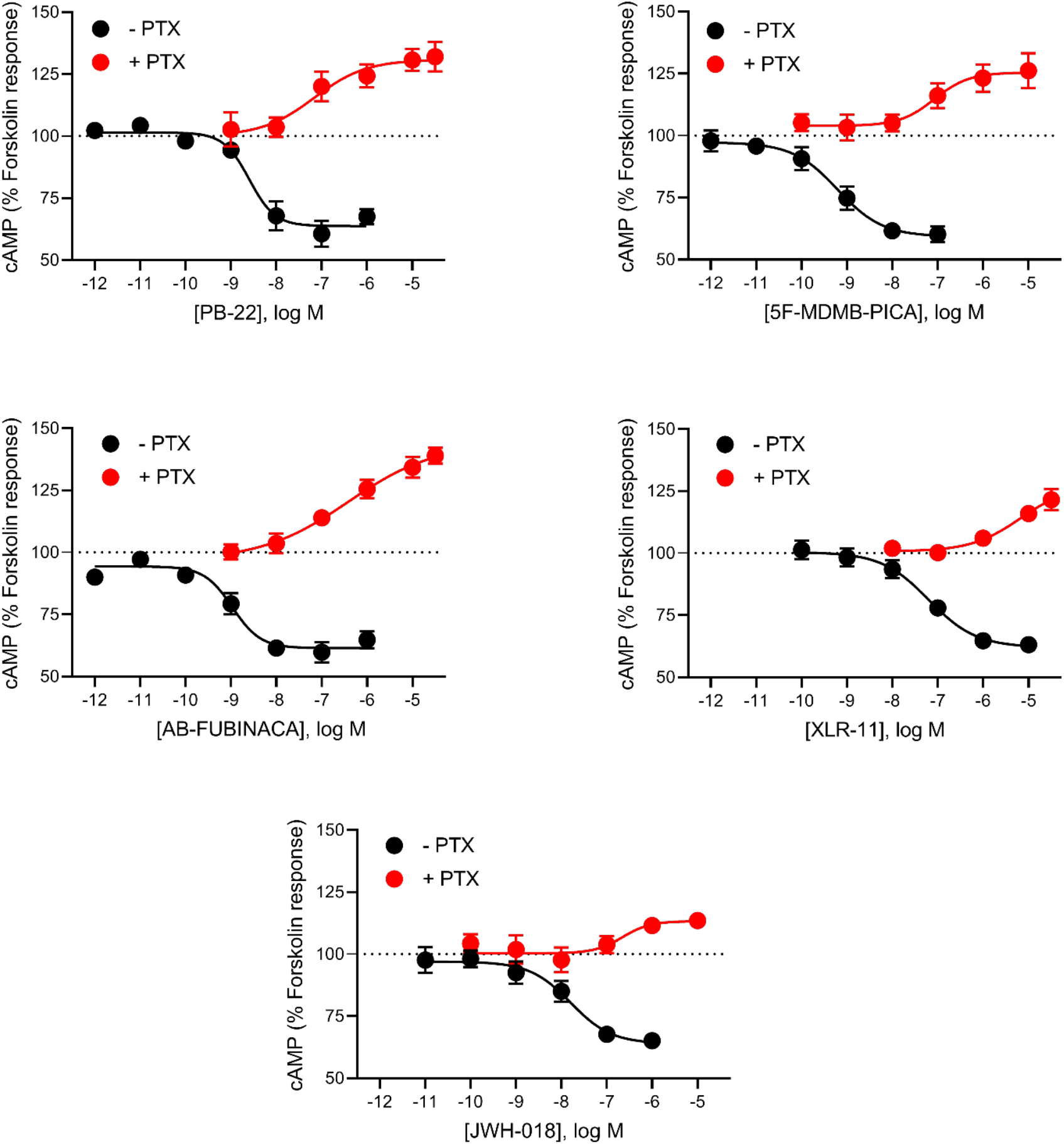
Concentration response curves for SCRAs-induced stimulation and inhibition of cAMP signalling. Concentration response relationship for five SCRAs (PB-22, 5F-MDMB-PICA, AB-FUBINACA, XLR-11, and JWH-018) for two signalling outputs of CB1 – stimulation and inhibition of cAMP levels following overnight treatment in the absence (− PTX, black), or presence (+ PTX, red) of PTX. Data were normalized to forskolin (100%) and vehicle (0%), and plotted as mean ± SEM for at least 5 independent experiments performed in duplicate. For some points, the error bars are shorter than the height of the symbol.

We then tested if the SCRA-induced observed stimulatory effects were mediated through CB1 receptors. Pre-treatment of HEK-CB1 with SR141716A (3 μM, 5 min), a potent and selective CB1 antagonist [30], largely prevented the subsequent SCRA (10 μM)-mediated stimulation of forskolin-induced cAMP response compared to the vehicle-treated cells (Figure 4, P < 0.05). Consistent with Gα_s_ CB1-specific responses of SCRAs, pre-treatment with SR141716A also blocked the inhibitory cAMP signalling induced by SCRAs (Supplementary Figure 1, P < 0.05).

**Figure 4.**
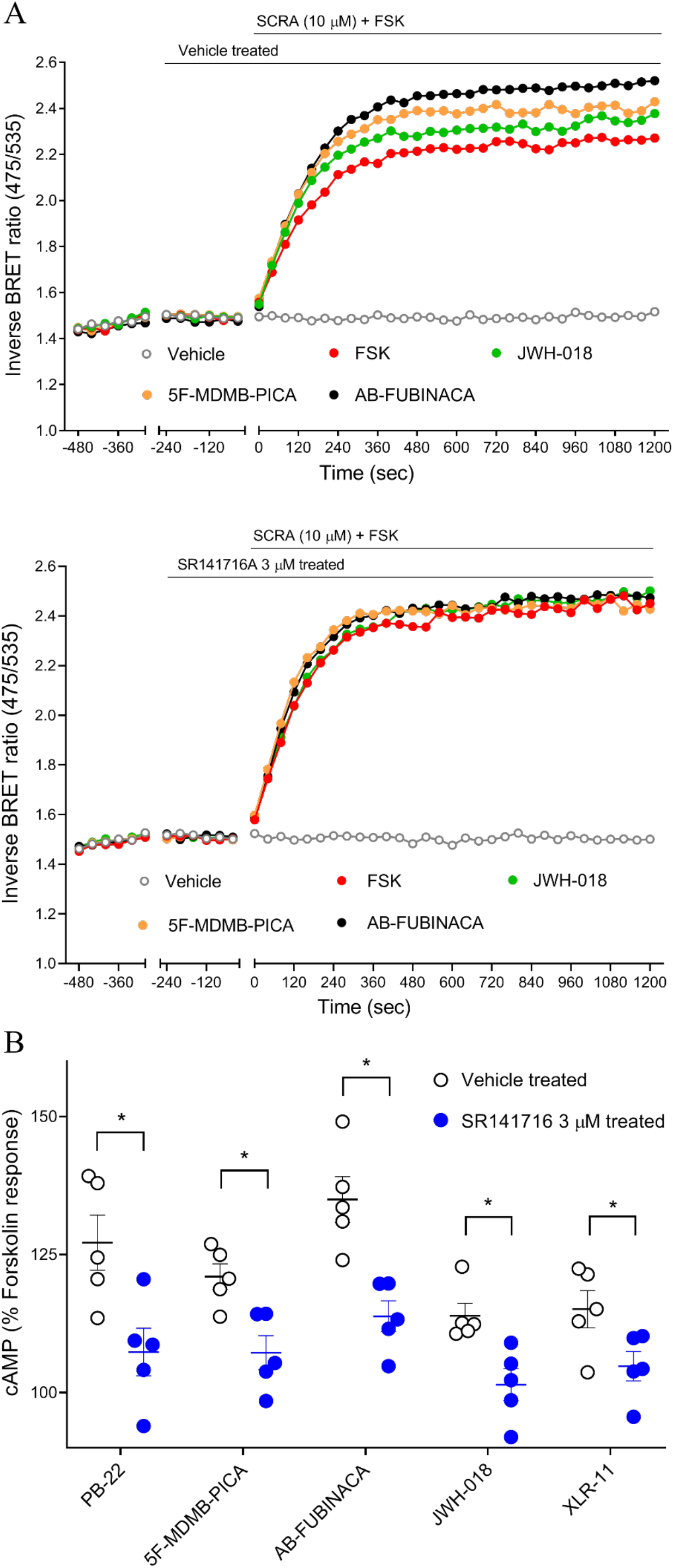
Effect of CB1 antagonist on the SCRA-mediated cAMP signalling peaks in HEK-CB1 cells. **A.** Traces from a representative experiment showing that SCRA (JWH-018, 5F-MDMB-PICA, and AB-FUBINACA) induced observed stimulatory effects were inhibited by SR141716A (CB1 antagonist, 3 μM) pre-treatment. **B.** Scatter dot plot representing SCRAs-mediated stimulation of forskolin-induced cAMP response in presence and absence of SR141716A 3 μM on HEK 293 cells expressing CB1. Within each set SCRAs (10 μM) were compared to SCRAs + SR141716 (Unpaired Student’s t-test, P < 0.05 marked with *). Data were normalized to forskolin (100%) and vehicle (0%), and plotted as mean ± SEM for at least 5 independent experiments performed in duplicate.

AB-CHMINACA has previously been reported to stimulate Gα_s_-like cAMP signalling pathway in a concentration dependent manner in HEK-CB1 cells [21]. Following PTX treatment, AB-CHMINACA increased cAMP levels above that of forskolin alone (Figure 5) in a concentration-dependent manner, with an increase of 86 ± 21% at 30 μM. However, in cells pre-treated with SR141716A (3 μM, 5 min), the stimulatory effects of AB-CHMINACA (10 μM) was only partially inhibited, in contrast to other SCRAs tested in the present study. To confirm that this response was at least in part non-CB1-mediated, AB-CHMINACA was tested in HEK 293 wild-type (WT) cells; in these cells, AB-CHMINACA (10 μM) also produced a small increase in forskolin-stimulated cAMP accumulation (Figure 5, 29 ± 10%), suggesting that some of these stimulatory effects were occurring via mechanism(s) unrelated to CB1 receptor activity.

**Figure 5.**
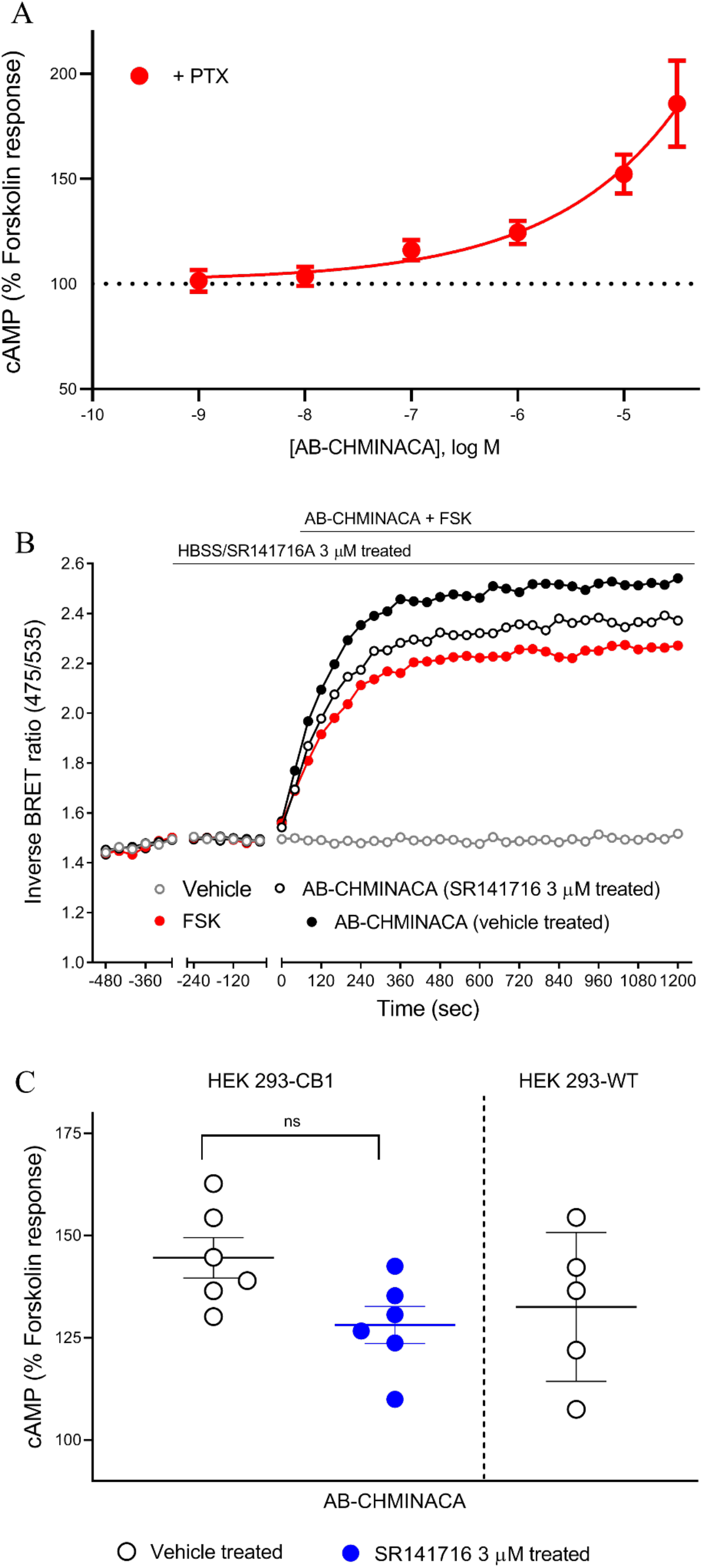
AB-CHMINACA does not modulate cAMP levels via CB1 receptors in HEK 293 cells. **A.** Treatment with AB-CHMINACA produced a concentration-dependent increase in forskolin-mediated cAMP production in HEK 293-CB1 in presence of PTX. **B.** Traces from a representative experiment showing that AB-CHMINACA (10 μM) induced observed stimulatory effects were only partially inhibited by SR141716A 3 μM. **C.** Scatter dot plot comparing AB-CHMINACA-mediated stimulation of forskolin-induced cAMP response in presence and absence of SR141716 3 μM in HEK 293-CB1 cells, and the data was not significantly different. AB-CHMINACA (10 μM) also modestly augmented forskolin-stimulated cAMP levels in HEK-WT cells (not containing CB1 receptors). Graphs show mean ± SEM for at least 5 independent experiments performed in duplicate.

## DISCUSSION

In this study, we set out to systematically characterize the ability of several SCRAs to activate Gα_s_ and Gα_i/o_ proteins by examining stimulation as well as inhibition of forskolin-induced cAMP accumulation in HEK cells stably expressing CB1. Assays of cAMP signalling revealed that the maximum concentration of SCRAs tested (10 μM), increased cAMP levels 12 to 45% above that produced by forskolin alone, while THC failed to increase cAMP levels, an observation consistent with the findings of Finlay et al. [17]. To further investigate the differential response of SCRA-induced activation and inhibition of cAMP production, we constructed the concentration response curves for the most prevalent group of SCRAs (JWH-018, PB-22, AB-FUBINACA, XLR-11, and 5F-MDMB-PICA); the rank order of potency of these SCRAs to stimulate Gα_s_-like cAMP signalling pathway was different from their activity at Gα_i/o_-pathway (inhibition of cAMP), suggesting that some of these drugs differentially regulate G protein coupling to CB1.

SCRA-mediated inhibition of cAMP has been extensively studied in cell models expressing cannabinoid receptors [21, 24]. Indeed, in some studies of CB1 signalling outputs, SCRAs have demonstrated Gα_s_-like phenotype [14, 15, 16, 17]. Initial experiments were conducted to determine whether traditional, endogenous, and synthetic cannabinoids stimulate Gα_s_ mediated stimulation of cAMP synthesis. Our results are consistent with the previous reports, showing the greater maximal effect of 2-AG at Gα_s_-like CB1 signalling compared to CP55940 and WIN55212-2 [17]. We found that 3 of the 16 SCRAs tested, AB-FUBINACA, PB-22, and AB-PINACA, activated Gα_s_-like CB1 signalling to more than 30% above the forskolin response. In a previous study using AB-CHMINACA, Costain and colleagues [21] showed similar increases in cAMP levels to that seen in this study without the need for FSK or PTX pre-treatment. Costain et al. [21], performed their assays on HEK293T cells transiently transfected with CB1. Transient transfection of CB1 may have led to a higher level of receptor expression than in our cells, and high levels of CB1 receptor expression is sufficient to result in a switch in cAMP signalling from Gα_i_-mediated (inhibitory) to Gα_s_-mediated (stimulatory) [17]. Furthermore, the HEK-293 “T” subclone used in the previous study harbors considerable genomic differences to the parental HEK 293 cell line used in the present study [31, 32], which may also contribute to altered cAMP responses (via different adenylyl cyclase isoforms). However, our data, together with that of Costain et al. [21] suggest potentially different receptor/effector coupling pathways in the presence of some SCRAs (AB-FUBINACA, PB-22, and AB-PINACA, AB-CHMINACA) compared to other CB1 ligands.

We further sought to investigate SCRA differential activation of distinctive G protein subsets—inhibition and stimulation of forskolin-mediated cAMP signalling. The relative ability of SCRAs to induce inhibition of cAMP production via Gα_i/o_ is very similar to that observed in previous studies in assays of membrane potential and [^35^S]GTPγS binding [12, 25, 33, 34]. The similar E_MAX_ observed for the SCRA-mediated activation of Gα_i/o_-CB1 signalling probably reflects receptor reserve for inhibition of cAMP accumulation in these cells, wherein maximal responses are elicited at less than maximal receptor occupancy because the system maximum is already achieved [12]. SCRA-induced stimulation of cAMP showed significant differences in E_MAX_ (Table 1), suggesting lower levels of receptor reserve for SCRAs coupled to Gα_s_ protein. This may (at least for the drugs with a lower E_MAX_) reflect an accurate representation of intrinsic efficacy of the ligands at this pathway [35]. The observed dynamic range of E_MAX_ for cannabinoids is consistent with CB1 having low coupling efficiency to both Gα_s_-pathway and β-arrestin-2 (as observed previously; [32], compared to that of Gα_i_-pathway [17, 36, 37]. Future studies could examine the structure of SCRA-bound CB1-Gα_s_ complexes, which might assist in explaining the observed cAMP signalling profiles. This is particularly interesting given that the interaction of SCRA MDMB-FUBINACA with the “toggle twin switch” in the CB1 binding pocket coupled to Gα_i_ was recently studied [38]. The rigid C-shape geometry of MDMB-FUBINACA along with the strong pi-pi interaction of its indazole ring with “toggle twin switch” residues, might help distinguish the high efficacy agonist activity of SCRA from partial agonists like THC lacking “toggle twin switch” interaction [38]. Promiscuous coupling to both Gα_i_ and Gα_s_ has been reported for multiple GPCRs (e.g. β_2_-adrenergic receptor) [39], while some receptors couple pre-dominantly to one G protein subtype (e.g. μ-opioid receptor coupling to the Gα_i/o_ family, [40]). The potential of cannabinoids to differentially activate one signalling cascade over another (functional selectivity, [41]) may aid the development of new therapeutic compounds with reduced psychoactive effects; a research domain that has attracted much recent interest [42].

Considering the adverse effects associated with SCRA use, it is important to continue characterizing the pharmacological profile of these compounds in order to understand the mechanisms driving their toxicity [43, 44]. Although this study does not identify which pathway contributes to the toxic effects observed following SCRA consumption, our data do provide valuable insights into SCRA-mediated stimulation and inhibition of cAMP signalling *in vitro*. Previous studies have shown that JWH-018- AM-2201-, 5F-AB-PINACA-, and CUMYL-4CN-BINACA-induced seizures are CB1-mediated in mice, which might explain some of the toxicity experienced by recreational users of these drugs [43, 44, 45, 46, 47, 48, 49]. Our data shows that SCRA-induced cAMP increase was abolished after SR141716A treatment, supporting the hypothesis that SCRAs Gα_s_-like effects were mediated through CB1 receptor. All the SCRAs tested in this study exhibited greater potency at Gα_i_- than Gα_s_-like pathways, and the efficacies of these SCRAs have previously been measured in response to Gα_i_-mediated activation of GIRK channel in AtT20-CB1 cells [12]. The rank order of SCRA efficacy based on selectivity for Gα_i_-GIRK signalling was found to be 5F-MDMB-PICA > XLR-11 > AB-FUBINACA > PB-22 ≈ JWH-018 [12]. 5F-MDMB-PICA showed the highest efficacy for modulation of K channel activity via Gα_i_-pathway in the former study, in contrast to the intermediate efficacy of 5F-MDMB-PICA to stimulate the Gα_s_-like cAMP signalling pathway in the present study. AB-FUBINACA exhibited greater efficacy for the Gα_s_-pathway compared to its Gα_i_-mediated activity profile in the membrane potential assay [12]. Evaluating the differences in G protein preference between SCRAs may be an important part of understanding the apparent differences in effect between these drugs in humans.

Our study showed that SCRAs have significantly different pharmacological profiles (maximal activities and potencies) for the activation of CB1-G protein-stimulation and -inhibition of forskolin-mediated cAMP signalling. Although it is speculated that the adverse effects of SCRAs are mediated by CB1 [49, 50], based on the results presented here we wonder how the differential responses of SCRAs are related to the physiological effects resulting from the activation of each intracellular pathway, and if these may be correlated with the *in vivo* toxicity of SCRAs. The unique toxicological profile of SCRAs may result from a combination of factors; pharmacokinetic differences, activity at both cannabinoid and non-cannabinoid targets, pharmacological activity of metabolites and thermolytic degradants [25, 37, 51, 52]. These findings may provide a starting point to help predict the pharmacological characteristics of SCRAs that demonstrate differential activation of Gα_i_ versus Gα_s_ coupling to CB1.

## Abbreviations

SCRA: synthetic cannabinoid receptor agonists
NPS: novel psychoactive substances
CB1: cannabinoid receptor type 1
HEK-CB1: Human Embryonic Kidney cells stably transfected with HA tagged human CB1 receptors
HA: haemagglutinin
AC: adenylyl cyclase
PTX: pertussis toxin
BRET: bioluminescence resonance energy transfer
CAMYEL: cAMP sensor YFP-Epac-Rluc
PEI: polyethylenimine
GIRK: G protein-gated inwardly rectifying K^+^ channel
2-AG: 2-arachidonoyl glycerol
THC: Δ^9^-tetrahydrocannabinol

## Acknowledgements

This work was supported by NHMRC Project Grant 1107088 awarded to M.K., and M.C. S.S. was supported by a Macquarie University Research Excellence Scholarship. The authors thank Dr Natasha Grimsey (Auckland University) for helpful advice on establishing the CAMYEL assay.

## Author contributions

S.S. designed and performed experiments, analysed the data and wrote the manuscript. S.D.B. synthesized SCRAs supervised by M.K. M.S. and C.B. advised on the CAMYEL assay and data analysis. M.C. provided critical feedback and helped shape the research, analysis, and manuscript. All authors reviewed and edited the manuscript.

---

**Supplementary Figure 1.**
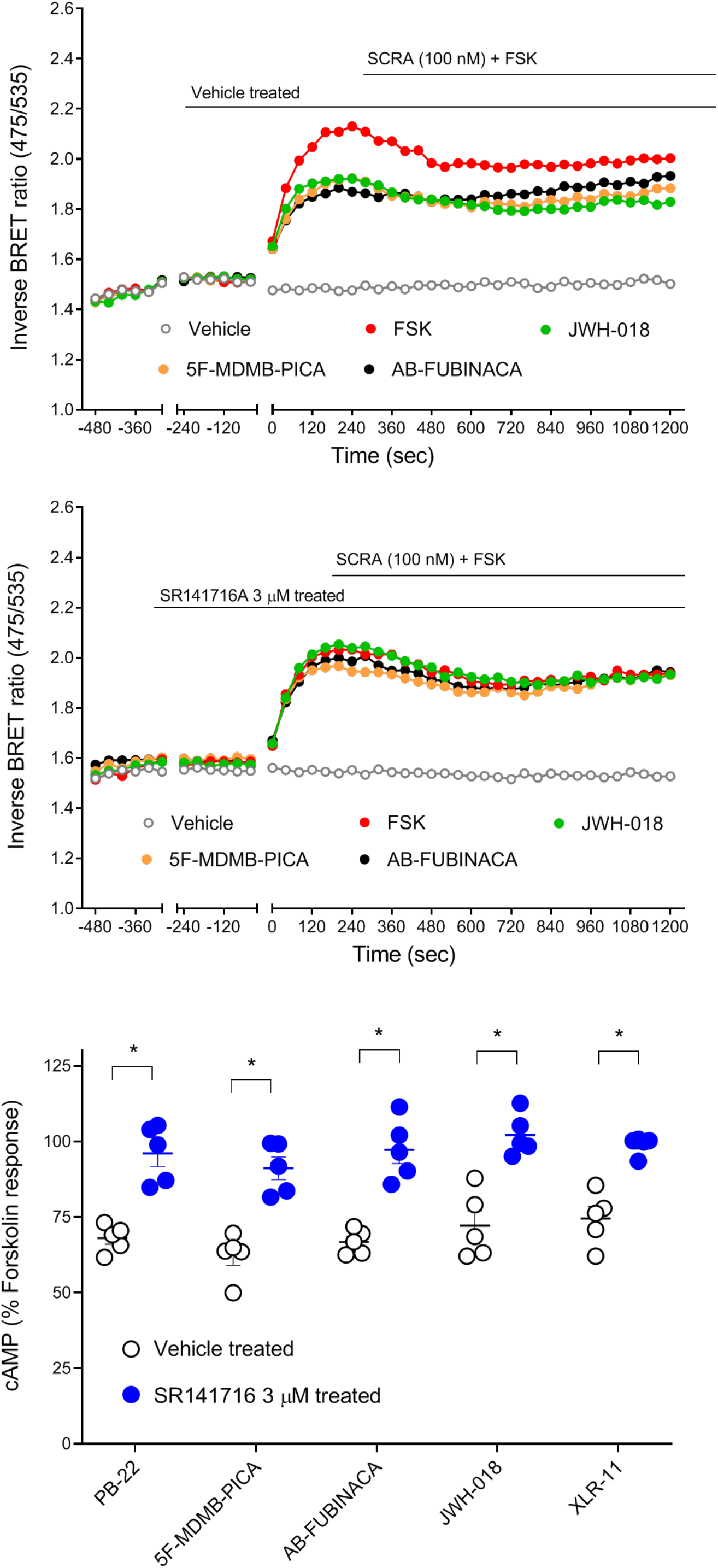
Effect of CB1 antagonist on the SCRA induced inhibition of cAMP signalling. **A.** Traces from a representative experiment showing that SCRA (JWH-018, 5F-MDMB-PICA, and AB-FUBINACA) induced inhibitory effects were completely blocked by SR141716A (CB1 antagonist, 3 μM) pre-treatment. **B.** Scatter dot plot representing SCRAs-mediated inhibition of forskolin-induced cAMP response in presence and absence of SR141716A 3 μM on HEK 293 cells expressing CB1. Within each set SCRAs (100 nM) were compared to SCRAs + SR141716. Data were normalized to forskolin (100%) and vehicle (0%), and plotted as mean ± SEM for at least 5 independent experiments performed in duplicate.

## References

1. Brandt SD, King LA, Evans‐Brown M. The new drug phenomenon. Drug Test Anal. 2014; 6: 587–597.

2. Wiley JL, Marusich JA, Lefever TW, et al. AB-CHMINACA, AB-PINACA, and FUBIMINA: Affinity and Potency of Novel Synthetic Cannabinoids in Producing Δ9-Tetrahydrocannabinol–Like Effects in Mice. J Pharmacol Exp Ther. 2015; 354:328–339.

3. Adams AJ, Banister SD, Irizarry L, Trecki J, Schwartz M, Gerona R. “Zombie” outbreak caused by the synthetic cannabinoid AMB-FUBINACA in New York. N Engl J Med. 2017; 376: 235–242.

4. Trecki J, Gerona RR, Schwartz MD. Synthetic cannabinoid-related illnesses and deaths. N Engl J Med. 2015; 373: 103–107.

5. Banister SD, Connor M. The chemistry and pharmacology of synthetic cannabinoid receptor agonists as new psychoactive substances: origins. Handb Exp Pharmacol. 2018; 252: 165–190.

6. Auwärter V, Dresen S, Weinmann W, Müller M, Pütz M, Ferreirós N. ‘Spice’and other herbal blends: harmless incense or cannabinoid designer drugs? J Mass Spectrom. 2009; 44: 832–837.

7. Thakur GA, Tichkule R, Bajaj S, Makriyannis A. Latest advances in cannabinoid receptor agonists. Expert Opin Ther Pat. 2009; 19: 1647–1673.

8. Aghazadeh Tabrizi M, Baraldi PG, Borea PA, Varani K. Medicinal chemistry, pharmacology, and potential therapeutic benefits of cannabinoid CB2 receptor agonists. Chem Rev. 2016; 116: 519–560.

9. United Nations Office on Drug and Crime. Global overview of drug demand and supply. United Nations 2019.

10. Alexander SP, Kelly E, Marrion NV, et al. The concise guide to pharmacology 2017/18: overview. Br J Pharmacol. 2017; 174: S1–S16.

11. Huestis MA, Gorelick DA, Heishman SJ, et al. Blockade of effects of smoked marijuana by the CB1-selective cannabinoid receptor antagonist SR141716. Arch Gen Psychiatry. 2001; 58: 322–328.

12. Sachdev S, Vemuri K, Banister S, et al. In vitro determination of the CB1 efficacy of illicit synthetic cannabinoids. Br J Pharmacol. 2019. In Press

13. Mallipeddi S, Janero DR, Zvonok N, Makriyannis A. Functional selectivity at G-protein coupled receptors: Advancing cannabinoid receptors as drug targets. Biochem Pharmacol. 2017;128: 1–11.

14. Glass M, Felder CC. Concurrent stimulation of cannabinoid CB1 and dopamine D2 receptors augments cAMP accumulation in striatal neurons: evidence for a Gs linkage to the CB1 receptor. J Neurosci. 1997; 17: 5327–5333.

15. Bonhaus D, Chang L, Kwan J, Martin G. Dual activation and inhibition of adenylyl cyclase by cannabinoid receptor agonists: evidence for agonist-specific trafficking of intracellular responses. J Pharmacol Exp Ther. 1998; 287: 884–888.

16. Scotter E, Goodfellow C, Graham E, Dragunow M, Glass M. Neuroprotective potential of CB1 receptor agonists in an in vitro model of Huntington’s disease. Br J Pharmacol. 2010; 160: 747–761.

17. Finlay DB, Cawston EE, Grimsey NL, et al. Gαs signalling of the CB1 receptor and the influence of receptor number. Br J Pharmacol. 2017; 174: 2545–2562

18. Laprairie RB, Bagher AM, Kelly ME, Dupré DJ, Denovan-Wright EM. Type 1 cannabinoid receptor ligands display functional selectivity in a cell culture model of striatal medium spiny projection neurons. J Biol Chem. 2014; 289: 24845–24862.

19. Lauckner JE, Hille B, Mackie K. The cannabinoid agonist WIN55, 212-2 increases intracellular calcium via CB1 receptor coupling to Gq/11 G proteins. Proceedings of the National Academy of Sciences. 2005;102(52):19144–19149.

20. Mukhopadhyay S, Howlett AC. Chemically distinct ligands promote differential CB1 cannabinoid receptor-Gi protein interactions. Mol Pharmacol. 2005; 67: 2016–2024.

21. Costain WJ, Rasquinha I, Comas T, et al. Analysis of the pharmacological properties of JWH-122 isomers and THJ-2201, RCS-4 and AB-CHMINACA in HEK293T cells and hippocampal neurons. Eur J Pharmacol. 2018; 823: 96–104.

22. Finlay DB, Manning JJ, Ibsen MS, et al. (2019). Do toxic synthetic cannabinoid receptor agonists have signature in vitro activity profiles ? A case study of AMB-FUBINACA. ACS Chemical Neuroscience, In Press

23. Schoeder CT, Hess C, Madea B, Meiler J, Müller CE. Pharmacological evaluation of new constituents of “Spice”: synthetic cannabinoids based on indole, indazole, benzimidazole and carbazole scaffolds. Forensic Toxicol. 2018; 36: 385–403.

24. Hess C, Schoeder CT, Pillaiyar T, Madea B, Müller CE. Pharmacological evaluation of synthetic cannabinoids identified as constituents of spice. Forensic Toxicol. 2016; 34: 329–343.

25. Wiley JL, Lefever TW, Marusich JA, et al. Evaluation of first generation synthetic cannabinoids on binding at non-cannabinoid receptors and in a battery of in vivo assays in mice. Neuropharmacology. 2016; 110: 143–153.

26. Knapman A, Abogadie F, McIntrye P, Connor M. A Real-Time, Fluorescence-Based Assay for Measuring μ-Opioid Receptor Modulation of Adenylyl Cyclase Activity in Chinese Hamster Ovary Cells. J Biomol Screen. 2014; 19: 223–231.

27. Jiang LI, Collins J, Davis R, et al. Use of a cAMP BRET sensor to characterize a novel regulation of cAMP by the sphingosine 1-phosphate/G13 pathway. J Biol Chem. 2007; 282: 10576–10584.

28. Sachdev S, Boyd R, Grimsey NL, Santiago M, Connor M. Brodifacoum does not modulate human cannabinoid receptor-mediated hyperpolarization of AtT20 cells or inhibition of adenylyl cyclase in HEK 293 cells. PeerJ. 2019; 7: e7733.

29. Cawston EE, Redmond WJ, Breen CM, Grimsey NL, Connor M, Glass M. Real‐time characterization of cannabinoid receptor 1 (CB 1) allosteric modulators reveals novel mechanism of action. Br J Pharmacol. 2013; 170: 893–907.

30. Rinaldi-Carmona M, Barth F, Héaulme M, et al. SR141716A, a potent and selective antagonist of the brain cannabinoid receptor. FEBS Lett. 1994; 350: 240–244.

31. Lin Y-C, Boone M, Meuris L, et al. Genome dynamics of the human embryonic kidney 293 lineage in response to cell biology manipulations. Nat Comm. 2014; 5: 4767.

32. Ibsen MS, Finlay DB, Patel M, Javitch JA, Glass M, Grimsey NL. Cannabinoid CB1 and CB2 receptor-mediated arrestin translocation: species, subtype, and agonist-dependence. Front Pharmacol. 2019; 10: 350.

33. Banister SD, Longworth M, Kevin R, et al. Pharmacology of valinate and tert-leucinate synthetic cannabinoids 5F-AMBICA, 5F-AMB, 5F-ADB, AMB-FUBINACA, MDMB-FUBINACA, MDMB-CHMICA, and their analogues. ACS Chem Neurosci. 2016; 7: 1241–1254.

34. Gamage TF, Farquhar CE, Lefever TW, et al. Molecular and Behavioral Pharmacological Characterization of Abused Synthetic Cannabinoids MMB-and MDMB-FUBINACA, MN-18, NNEI, CUMYL-PICA, and 5-Fluoro-CUMYL-PICA. J Pharmacol Exp Ther. 2018; 365: 437–446.

35. Kenakin T. New concepts in pharmacological efficacy at 7 TM receptors: IUPHAR Review 2. Br J Pharmacol. 2013; 168: 554–575.

36. Wouters E, Walraed J, Banister SD, Stove CP. Insights into biased signaling at cannabinoid receptors: synthetic cannabinoid receptor agonists. Biochem Pharmacol. 2019; 169: 113623

37. Cannaert A, Storme J, Franz F, Auwärter V, Stove CP. Detection and activity profiling of synthetic cannabinoids and their metabolites with a newly developed bioassay. Anal Chem. 2016; 88: 11476–11485.

38. Kumar KK, Shalev-Benami M, Robertson MJ, et al. Structure of a signaling cannabinoid receptor 1-G protein complex. Cell. 2019; 176: 448–458.

39. Westfield GH, Rasmussen SG, Su M, et al. Structural flexibility of the Gαs α-helical domain in the β2-adrenoceptor Gs complex. Proc Natl Acad Sci. 2011; 108: 16086–16091.

40. Connor M, Christie MJ. Opioid receptor signalling mechanisms. Clin Exp Pharmacol Physiol. 1999; 26: 493–499.

41. Kenakin T. Functional selectivity and biased receptor signaling. J Pharmacol Exp Ther. 2011; 336: 296–302.

42. Wouters E, Walraed J, Robertson MJ, et al. Assessment of biased agonism amongst distinct synthetic cannabinoid receptor agonist scaffolds. ACS Pharmacol Transl Sci. 2019; In Press

43. Drummer OH, Gerostamoulos D, Woodford NW. Cannabis as a cause of death: A review. Forensic Sci Int. 2019; 298: 298–306.

44. Labay LM, Caruso JL, Gilson TP, et al. Synthetic cannabinoid drug use as a cause or contributory cause of death. Forensic Sci Int. 2016; 260: 31–39.

45. Funada M, Takebayashi-Ohsawa M. Synthetic cannabinoid AM2201 induces seizures: Involvement of cannabinoid CB1 receptors and glutamatergic transmission. Toxicol Appl Pharmacol. 2018; 338: 1–8.

46. Malyshevskaya O, Aritake K, Kaushik MK, et al. Natural (Δ 9-THC) and synthetic (JWH-018) cannabinoids induce seizures by acting through the cannabinoid CB 1 receptor. Sci Rep. 2017; 7: 10516.

47. Vigolo A, Ossato A, Trapella C, et al. Novel halogenated derivates of JWH-018: behavioral and binding studies in mice. Neuropharmacology. 2015; 95: 68–82.

48. Wilson CD, Tai S, Ewing L, et al. Convulsant effects of abused synthetic cannabinoids jwh-018 and 5f-ab-pinaca are mediated by agonist actions at cb1 receptors in mice. J Pharmacol Exp Ther. 2019; 368: 146–156.

49. Kevin RC, Anderson L, McGregor IS, et al. CUMYL-4CN-BINACA is an efficacious and potent pro-convulsant synthetic cannabinoid receptor agonist. Front Pharmacol. 2019;10.

50. Wallace MJ, Wiley JL, Martin BR, DeLorenzo RJ. Assessment of the role of CB1 receptors in cannabinoid anticonvulsant effects. Eur J Pharmacol. 2001; 428: 51–57.

51. Schindler CW, Gramling BR, Justinova Z, Thorndike EB, Baumann MH. Synthetic cannabinoids found in “spice” products alter body temperature and cardiovascular parameters in conscious male rats. Drug Alch Dep. 2017; 179: 387–394.

52. Thomas BF, Lefever TW, Cortes RA, et al. Thermolytic degradation of synthetic cannabinoids: chemical exposures and pharmacological consequences. J Pharmacol Exp Ther. 2017; 361: 162–171.

